## Supplementary material for "Machine learning models reveal environmental and genetic factors associated with the plant circadian clock"

| Gene ID | Name | meta2d Phase | meta2d Period | meta2d Q-value |
|---|---|---|---|---|
| AT1G01060 | *LHY* | 23.69 | 24.10 | 2.25E-06 |
| AT1G22770 | *GI* | 9.44 | 23.95 | 1.29E-05 |
| AT2G21070 | *FIO1* | 15.02 | 18.07 | 0.07 |
| AT2G25930 | *ELF3* | 15.92 | 22.85 | 1.54E-05 |
| AT2G31870 | *TEJ* | 13.95 | 23.09 | 0.004 |
| AT2G46790 | *PRR9* | 5.83 | 24.17 | 0.0002 |
| AT2G46830 | *CCA1* | 0.88 | 24.40 | 1.80E-05 |
| AT3G20810 | *JMJD5* | 15.17 | 23.15 | 0.0006 |
| AT3G22380 | *TIC* | 15.85 | 22.29 | 0.005 |
| AT3G46640 | *PCL1* | 12.46 | 23.81 | 0.006 |
| AT4G39620 | *EMB2453* | 15.24 | 22.24 | 0.04 |
| AT5G02810 | *PRR7* | 7.48 | 24.17 | 1.34E-05 |
| AT5G42900 | *COR27* | 13.11 | 23.95 | 0.0001 |
| AT5G57360 | *ZTL* | 13.35 | 24.02 | 0.0005 |
| AT5G59570 | *BOA* | 14.98 | 24.25 | 0.0007 |
| AT5G60100 | *PRR3* | 13.77 | 23.15 | 0.0002 |
| AT5G61380 | *TOC1* | 13.75 | 23.28 | 0.0001 |

**Supplementary Table 1:** List of 17 clock genes selected based on prior biological knowledge and the fact they are consistently expressed across the training data. Includes approximations for phase period length and rhythmicity Q-values as determined using MetaCycle[1] within the continuous-light (LL) time-course by *Romanowski et al.*[2]

| Model | *N* gene features | Parameters | Tuning scheme | Tuned MAE (mins) |
|---|---|---|---|---|
| ZeitZeiger | 89 | *sumabsv* (regularization): 5<br>N sparse principal-components: 2 | LOO cross-validation | 52.9 (± 42.0) |
| PLSR ensemble | 233 (out of 100 sets) | N latent variables: 7 | 5-fold cross-validation | 14.8 (± 14.5) |
| Taufisher | 22 (484 gene-pair features) | N principal-components: 3 | LOO cross-validation | 88.9 (± 63.1) |
| MolecularTimetable **(WSN)** | 189 | N phases: 144<br>Pearson's *r* threshold > 0.89<br>Variation threshold > 0.15<br>Within-study-normalization | Training data | 47.1 (± 63.2) |
| TimeSignatR **(WSN)** | 167 | *a* (regularization): : 0.25<br>λ (shrinkage): exp(-2.0)<br>Within-study-normalization | LOO cross-validation | 14.3 (± 11.4) |

**Supplementary Table 2:** List of previously published circadian time (CT) predictors trained to compare with ChronoGauge including ZeitZeiger[3], partial-least-squares-regression[4] (PLSR as an ensemble), Taufisher[5], MolecularTimetable[6] and TimeSignatR[7]. Includes *N* gene features used an an input, the parameters & normalization techniques selected, the methods used to find said parameters and the mean absolute-errors (MAEs) of the tuned models across training/cross-validation data. We note MolecularTimetable was tuned without cross-validation because cosine wave fitting may not fit appropriately across folds. **LOO: Leave-one-out, WSN: within-study-normalization**

| RNA-seq benchmark test | | | | | | |
|---|---|---|---|---|---|---|
| Model | MdAE (mins) | | MAE (mins) | | *r* | |
| | Non-corrected | Combat-seq | Non-corrected | Combat-seq | Non-corrected | Combat-seq |
| ChronoGauge (x100) | **20.6 (± 47.6)** | 27.5 (± 40.7) | 44.0 (± 47.6) | **43.2 (± 40.7)** | 0.990 | **0.992** |
| ZeitZeitger | 46.2 (± 69.2) | 32.9 (± 46.9) | 73.8 (± 69.2) | 48.9 (± 46.9) | 0.976 | 0.991 |
| Taufisher | 90.0 (± 68.0) | 60.0 (± 57.0) | 105.5 (± 68.0) | 92.1 (± 57.0) | 0.985 | 0.991 |
| PLSR (x100) | 38.4 (± 49.7) | 37.5 (± 39.4) | 59.8 (± 49.7) | 48.2 (± 39.4) | 0.985 | 0.991 |
| MolecularTimetable | 50.0 (± 47.0) | 60.0 (± 45.6) | 54.8 (± 47.0) | 60.3 (± 45.6) | 0.989 | 0.988 |
| TimeSignatR | 60.2 (± 49.2) | 45.5 (± 46.4) | 65 (± 49.2) | 56.0 (± 46.4) | 0.983 | 0.987 |

**Supplementary Table 3:** Full list of evaluation metrics for each model's circadian time (CT) predictions in a hold-out RNA-seq set[8–13] (*N* samples = 58) using both non-corrected and Combat-seq[14] corrected expression values for training and testing. Includes median-absolute-error (MdAE), mean-absolute-error (MAE) and Pearson correlation coefficient (*r*). Paratheses denote standard-deviation of absolute-errors. Top scores for each metric listed in **bold**.

**x100**: ensemble of 100 sub-predictors, **PLSR**: Partial-least-squares-regression

| ATH1 microarray benchmark test set | | | | | | |
|---|---|---|---|---|---|---|
| Model | MdAE (mins) | | MAE (mins) | | *r* | |
| | Non-corrected | Combat-seq | Non-corrected | Combat-seq | Non-corrected | Combat-seq |
| ChronoGauge (x100) | **46.1 (± 50.2)** | 53.5 (± 57.0) | **61.8 (± 50.2)** | 73.0 (± 57.0) | **0.984** | 0.982 |
| ZeitZeitger | 77.9 (± 103.7) | 118.1 (± 127.4) | 123.8 (± 103.7) | 155.9 (± 127.4) | 0.932 | 0.564 |
| Taufisher | 120.0 (± 93.6) | 60.0 (± 74.0) | 116.7 (± 93.6) | 97.8 (± 74.0) | 0.946 | 0.983 |
| PLSR (x100) | 83.2 (± 77.6) | 98.1 (± 107.9) | 105.3 (± 77.6) | 124.0 (± 107.9) | 0.965 | 0.942 |
| MolecularTimetable | 70.0 (± 44.3) | 60.0 (± 50.9) | 64.7 (± 44.3) | 71.8 (± 50.9) | 0.984 | 0.980 |
| TimeSignatR | 75.3 (± 67.9) | 69.8 (± 64.6) | 82.3 (± 67.9) | 77.9 (± 64.6) | 0.971 | 0.903 |

**Supplementary Table 4:** Full list of evaluation metrics for each model's circadian time (CT) predictions in a hold-out ATH1 microarray set[15–18] (*N* samples = 73) using both non-corrected and Combat-seq[14] corrected expression values for training. Microarray test expression values were not batch corrected. Includes median-absolute-error (MdAE), mean-absolute-error (MAE) and Pearson correlation coefficient (*r*). Paratheses denote standard-deviation of absolute-errors. Top scores for each metric listed in **bold**.

**x100**: ensemble of 100 sub-predictors, **PLSR**: Partial-least-squares-regression

| AraGene microarray benchmark test set | | | | | | |
|---|---|---|---|---|---|---|
| Model | MdAE (mins) | | MAE (mins) | | *r* | |
| | Non-corrected | Combat-seq | Non-corrected | Combat-seq | Non-corrected | Combat-seq |
| ChronoGauge (x100) | **74.8 (± 68.1)** | 96.9 (± 88.8) | **89.7 (± 68.1)** | 11.7.4 (± 88.8) | **0.983** | 0.974 |
| ZeitZeitger | 122.0 (± 156.6) | 285.9 (± 202.2) | 160.3 (± 156.6) | 143.2 (± 110.8) | 0.911 | 0.629 |
| Taufisher | 120.0 (± 113.8) | 120.0 (± 92.7) | 156.7 (± 113.8) | 150 (± 92.7) | 0.937 | 0.963 |
| PLSR (x100) | 85.5 (± 99.7) | 103.5 (± 91.9) | 128.6 (± 99.7) | 128.6 (± 91.9) | 0.957 | 0.964 |
| MolecularTimetable | 100.0 (± 78.8) | 110.0 (± 71.6) | 118.3 (± 78.8) | 122.9 (± 71.6) | 0.974 | 0.977 |
| TimeSignatR | 116.0 (± 86.4) | 104.2 (± 79.0) | 126.2 (± 86.4) | 119.1 (± 79.0) | 0.959 | 0.949 |

**Supplementary Table 5:** Full list of evaluation metrics for each model's circadian time (CT) predictions in a hold-out AraGene microarray set[19] (*N* samples = 72) using both non-corrected and Combat-seq corrected expression values for training. Microarray test expression values were not batch corrected. Includes median-absolute-error (MdAE), mean-absolute-error (MAE) and Pearson correlation coefficient (*r*). Paratheses denote standard-deviation of absolute-errors. Top scores for each metric listed in **bold**.

**x100**: ensemble of 100 sub-predictors, **PLSR**: Partial-least-squares-regression

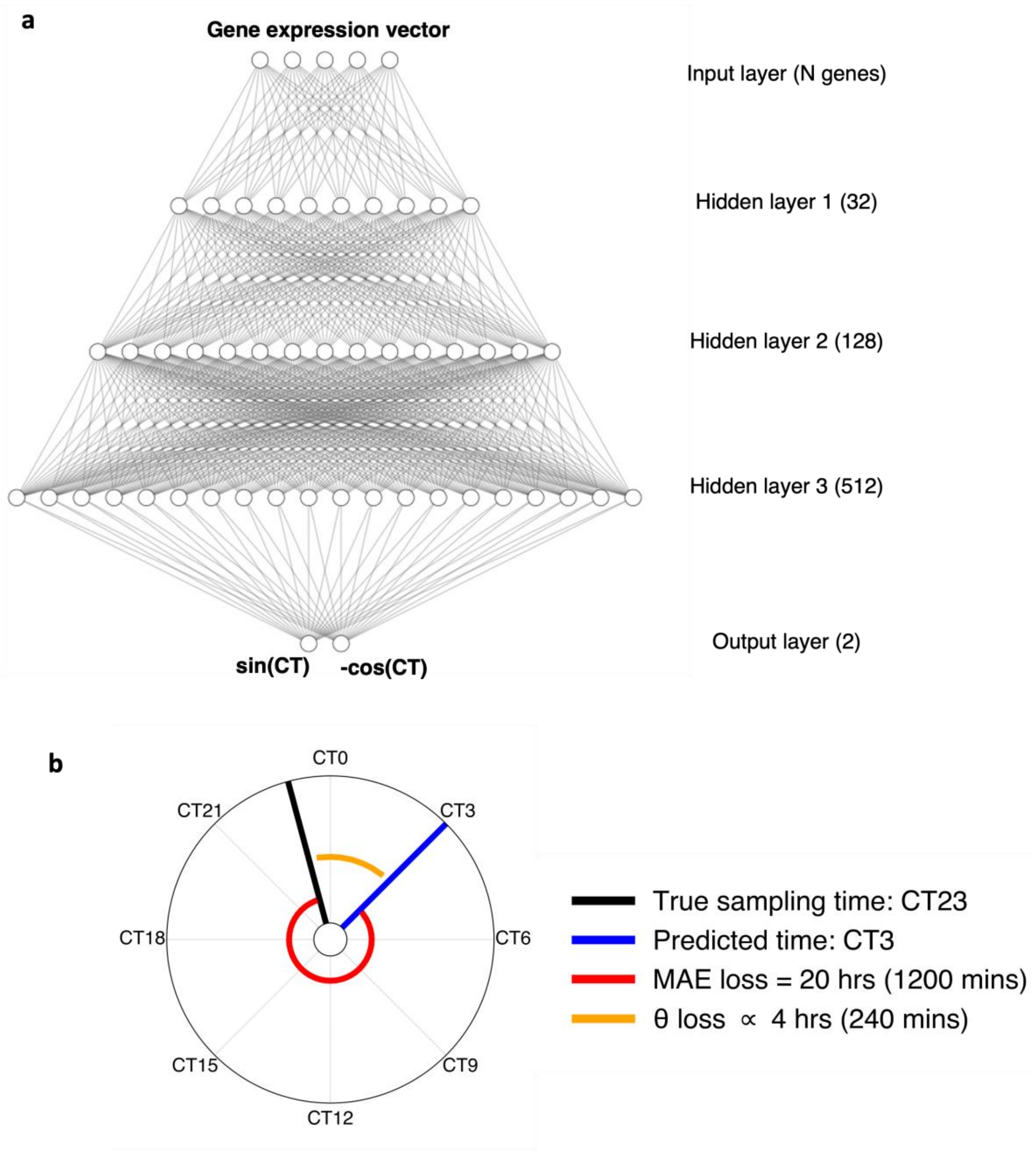


**Supplementary Figure 1:** Neural network architecture used by ChronoGauge based on a previously published model by *Gardiner et al*.[20]. **a** Multi-layer-perceotron (MLP) composed of 3 hidden layers that outputs the circadian time (CT) as sine and cosine values. **b** Justification of using θ (angle; orange) between true CT at CT23 (black) and predicted CT at CT3 as a loss function compared with mean-absolute-error (MAE) (red). A MAE loss does not consider the circular nature of the CT, while the θ loss does.

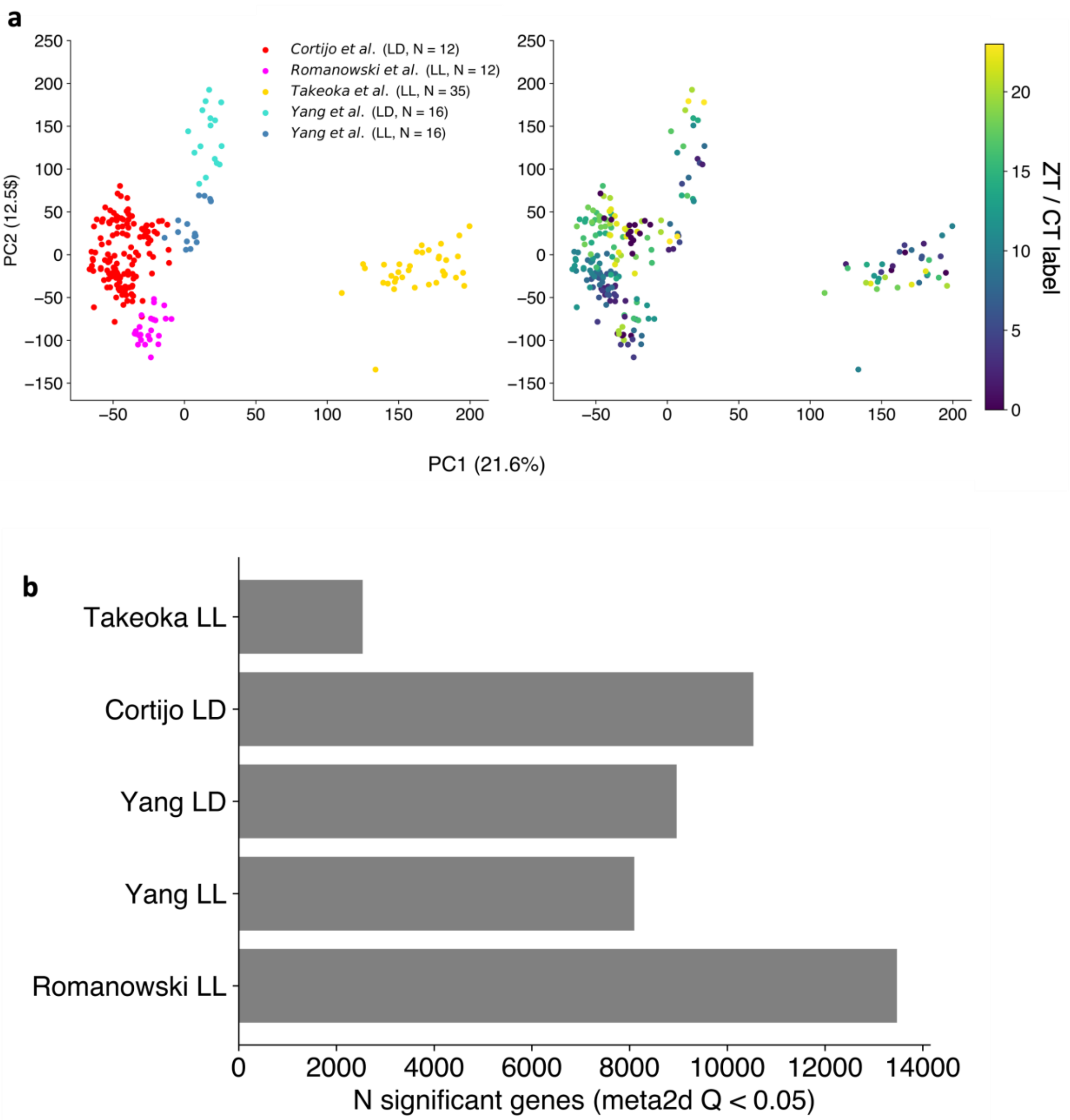


**Supplementary Figure 2:** Appraisal of the suitability of time-course RNA-seq datasets[2,21–23] harvested under either continuous-light (LL) or a light-dark cycle (LD) for training. **a** variation of proposed test samples labelled by experimental group (left) and circadian time (CT) or zeitgeber time (ZT). **b** Number of genes called as significantly rhythmic across training datasets based on results from MetaCycle[1] (meta2d Q < 0.05).

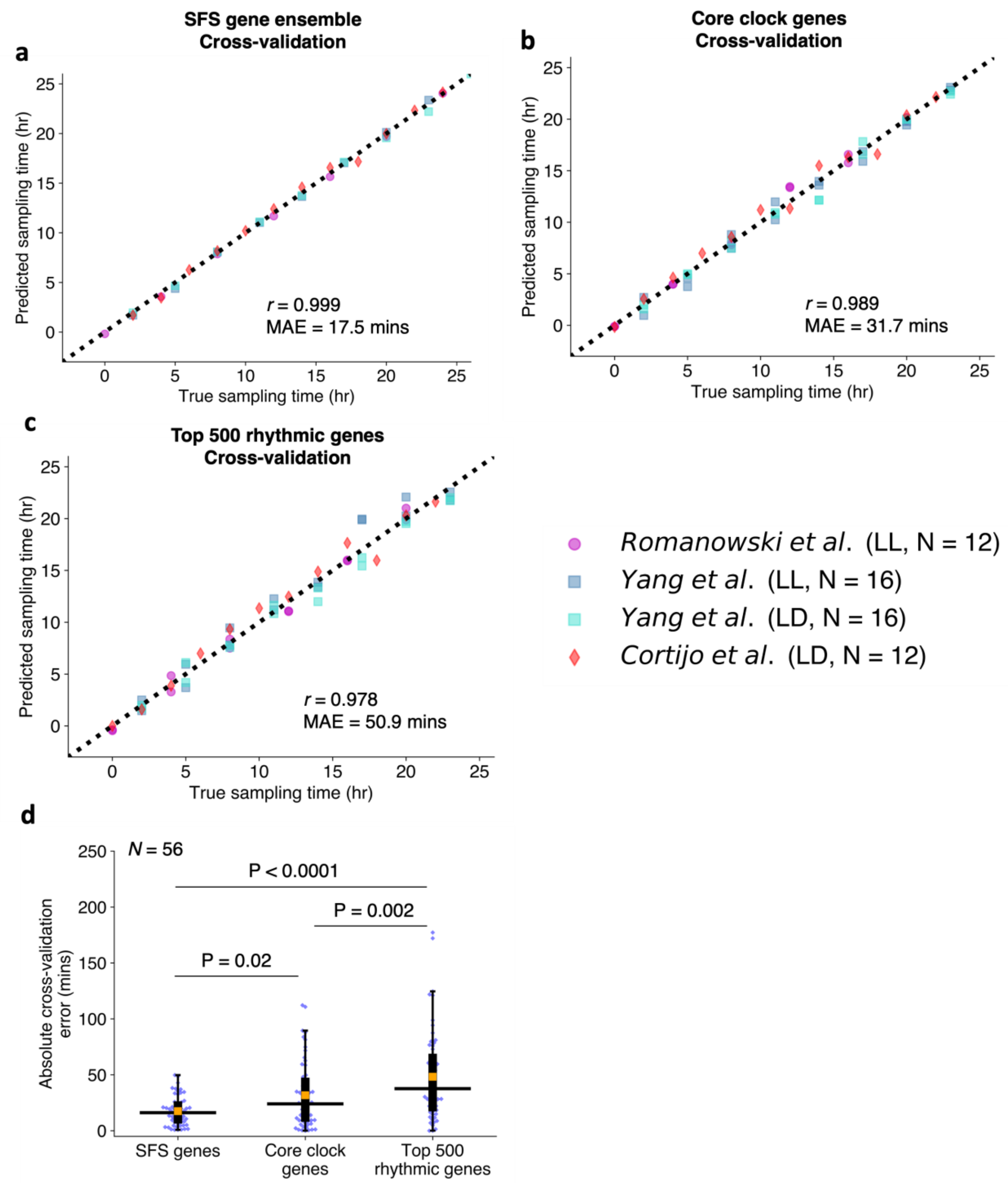


**Supplementary Figure 3:** Evaluation of tuned neural network (NN) models across 5-fold cross-validation for model selection using a combination of four separate time-course experiments[2,22,23] as a training dataset. Pearson correlation coefficients (*r)* and mean-absolute-erorrs (MAE) of circadian time (CT) estimates made for **a** using the sequentially selected feature (SFS) ensemble of 100 sub-predictors, **b** a NN trained using 17 core clock genes as features and **c** a NN trained using the top 500 rythmic genes as determined by meta2d Q-values as features. **d** Comparison of CT absolute-errors (mins) across 5-fold cross-validation including median-absolute-errors (MdAEs; horizontal black lines) and MAEs (orange boxes). MdAEs compared across each NN-model using a two-tailed Wilcoxon signed-rank test with a Bonferroni adjusted P-values. Experiments harvested under either continuous-light (LL) or diurnal/light-dark (LD) conditions are listed.

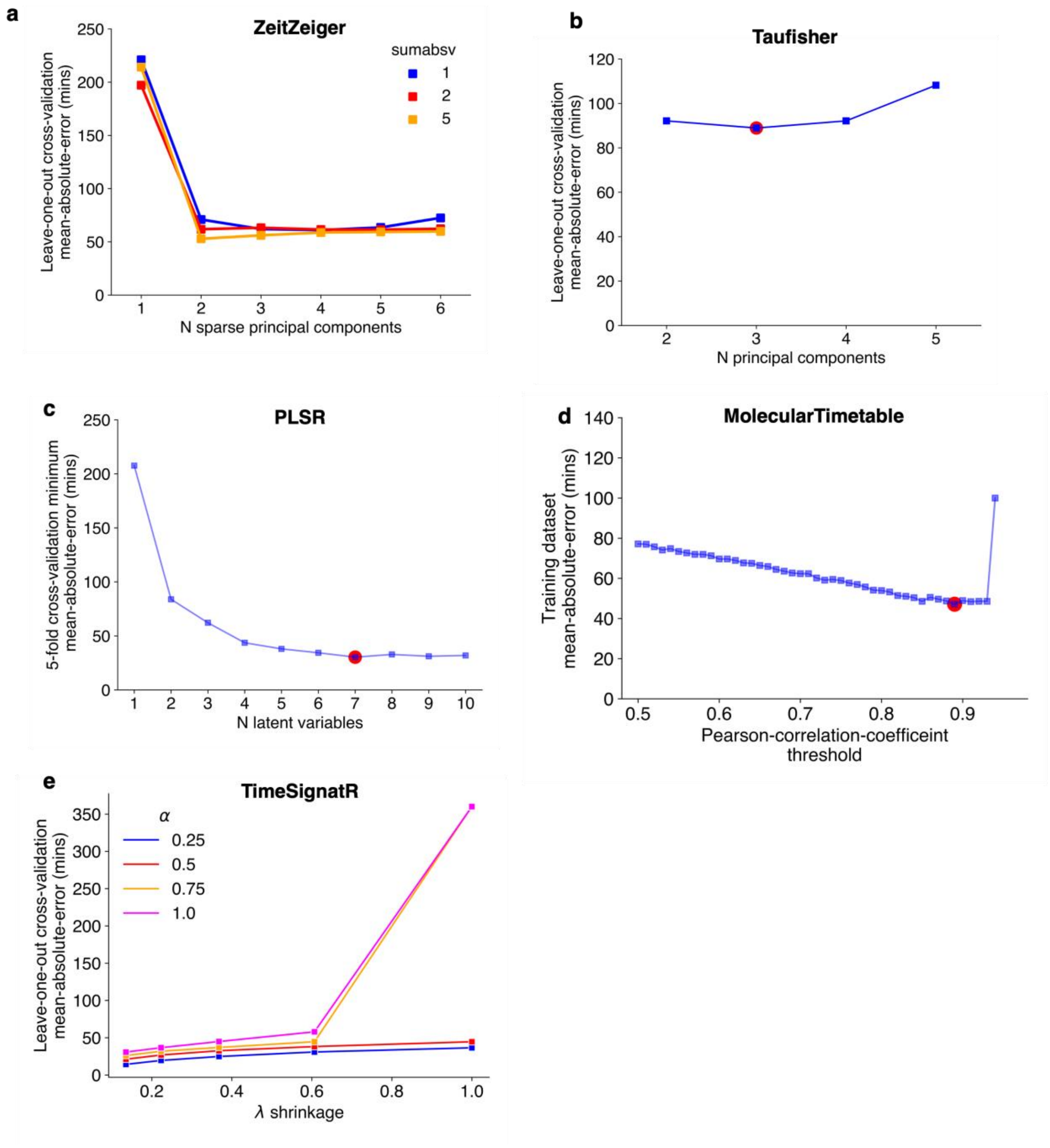


**Supplemental Figure 4:** Hyperparameter optimization of different circadian time (CT) prediction models using mean-absolute-error (MAE) across the same four time-course experiments[2,22,23] used to train ChronoGauge. **a,** Optimization of sumabsv (regularization factor) and *N* sparse principal components used for ZeitZeiger[3] using leave-one-out (LOO) cross-validation. **b,** Identification of optimal (red circle) *N* principal components used for Taufisher[5] using LOO cross-validation. **c,** Identification of the optimal (red circle) *N* latent variables used by partial-least-squares-regression[4] (PLSR) based on the minimum 5-fold cross-validation MAE displayed across a recursive feature elimination of 500 - 5 genes features. **d**, Identification of the optimal (red circle) Pearson correlation coefficient threshold (*r*) used to select time-indicating genes for MolecularTimetable[6] across the training data. **e,** Identification of the optimal α (regularization factor) and γ (shrinkage factor) used for TimeSignatuR[7] using LOO cross-validation.

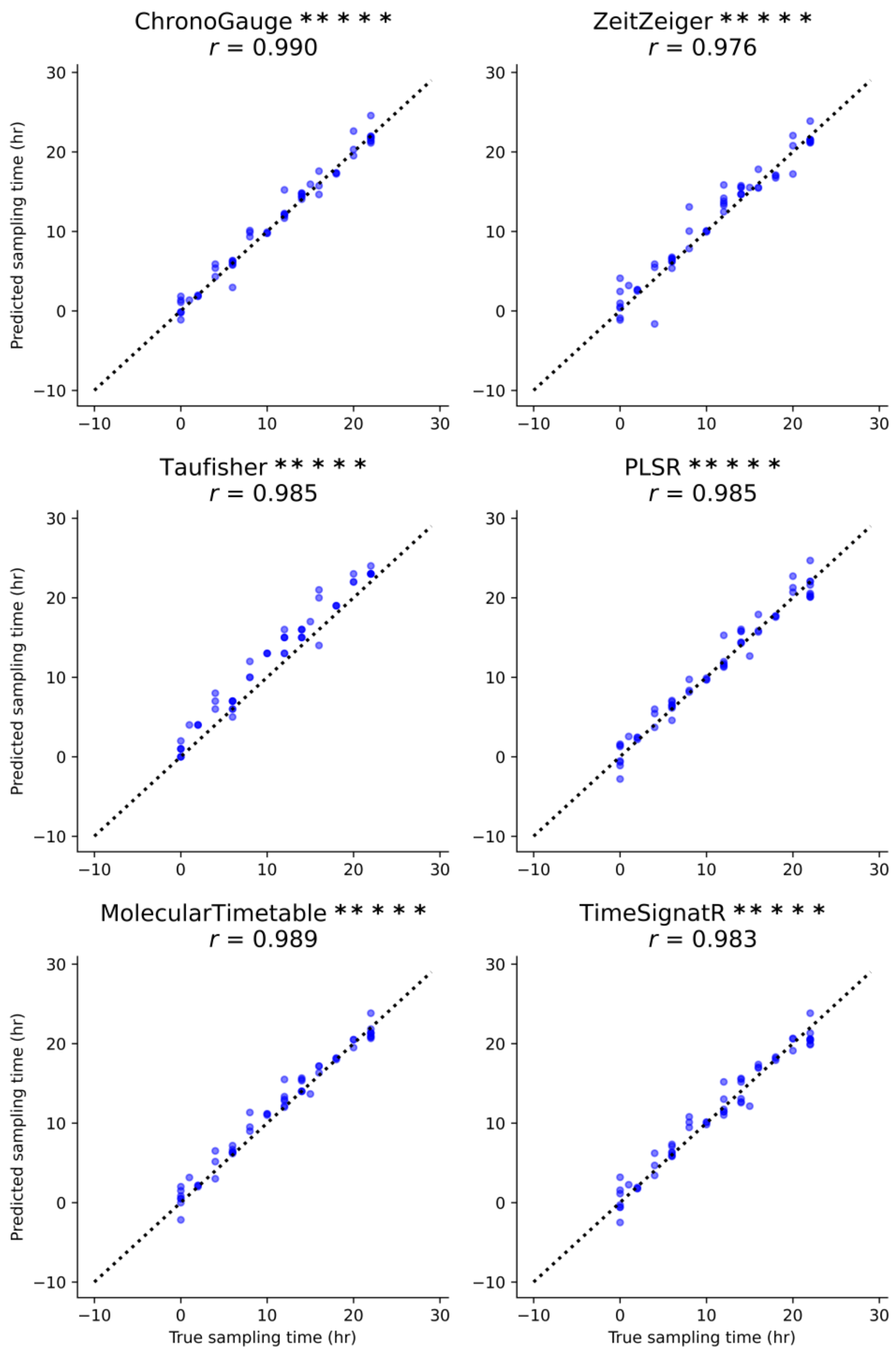


**Supplemental Figure 5:** Correlation of RNA-seq test samples[8–13] (*N* = 58) true harvesting time labels compared with the predicted circadian times (CT) across different models. CT predictions were adjusted to account for the 24-hour modulus. Pearson correlation coefficients (*r)* and P values shown. P values adjusted using Bonferroni method.

******* P < 0.00001**

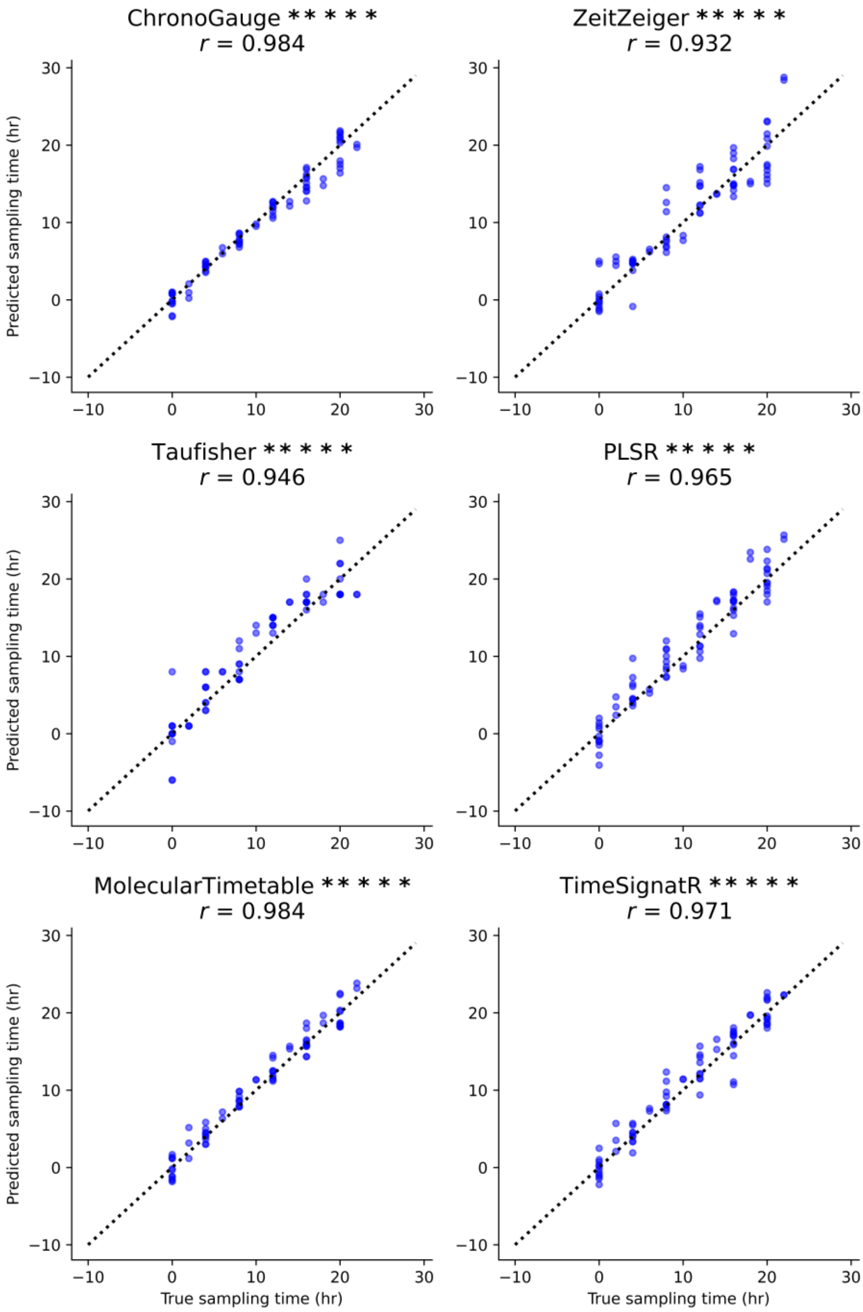


**Supplemental Figure 6:** Correlation of microarray ATH1 test samples[15–18] (*N* = 73) true harvesting time labels compared with the predicted circadian times (CT) across different models. CT predictions were adjusted to account for the 24-hour modulus. Pearson correlation coefficients (*r)* and P values shown. P values adjusted using Bonferroni method.

******* P < 0.00001**

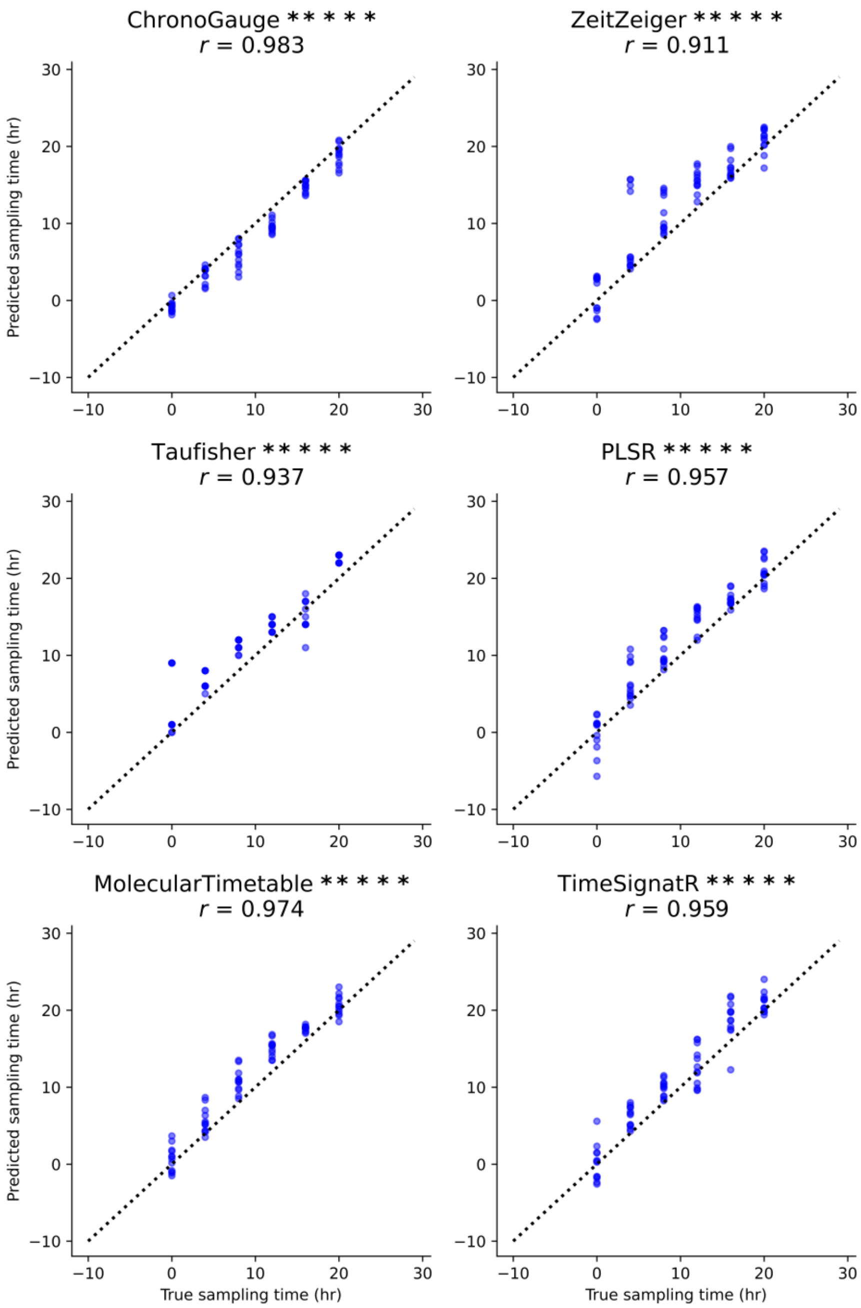


**Supplemental Figure 7:** Correlation of microarray AraGene test samples[19] ($N = 72$) true harvesting time labels compared with the predicted circadian times (CT) across different models. CT predictions were adjusted to account for the 24-hour modulus. Pearson correlation coefficients (*r)* and P values shown. P values adjusted using Bonferroni method.

******* P < 0.00001**

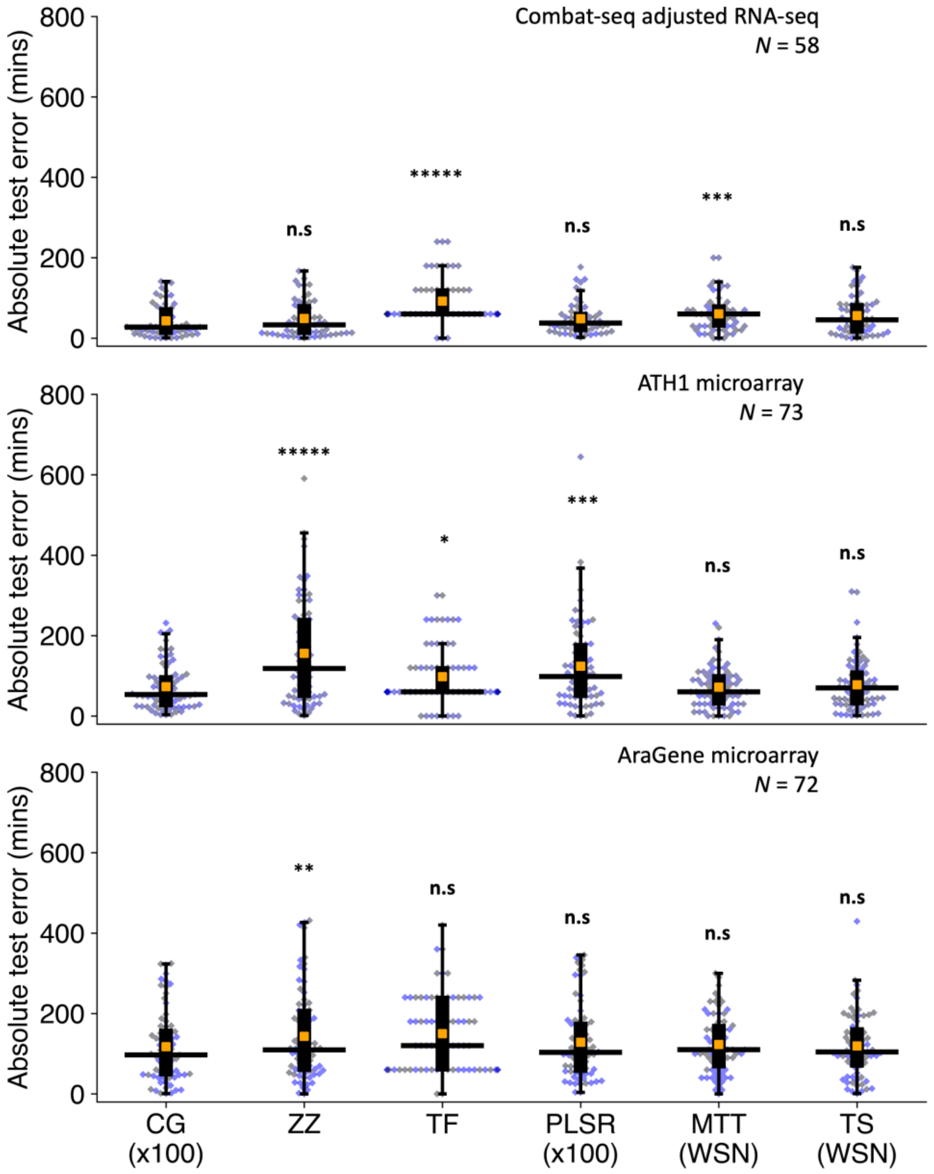


**Supplementary Figure 8:** Using Combat-seq adjusted RNA-seq expression as an input, comparison of hold-out test set absolute errors between ChronoGauge ensemble (CG x100) and competing circadian time (CT) prediction methods including ZeitZeiger[3] (ZZ), Taufisher[5] (TF), partial-least-squares-regression[4] ensemble (PLSR x100), MolecularTimetable[6] (MTT) and TimeSignatR[7] (TS) with boxplot including the median-absolute-error (MdAE; black line), mean-absolute-error (orange box) and the error for each sample (blue dot). Comparison includes a Combat-seq[14] adjusted RNA-seq set[8–13] (*N* samples = 58), an unadjusted ATH1 microarray set[15–18] (*N* samples = 73) and an unadjusted AraGene microarray set[19] (*N* samples = 72). Methods using within-study-normalization (WSN) listed. Microarray set signals were not adjusted. Bonferroni-adjusted P-values shown from a Wilcoxon signed-rank test.

**n.s** no significance**,** * P < 0.05, ** P < 0.01**,** *** P < 0.001**,** **** P < 0.0001**,** ***** P < 0.00001

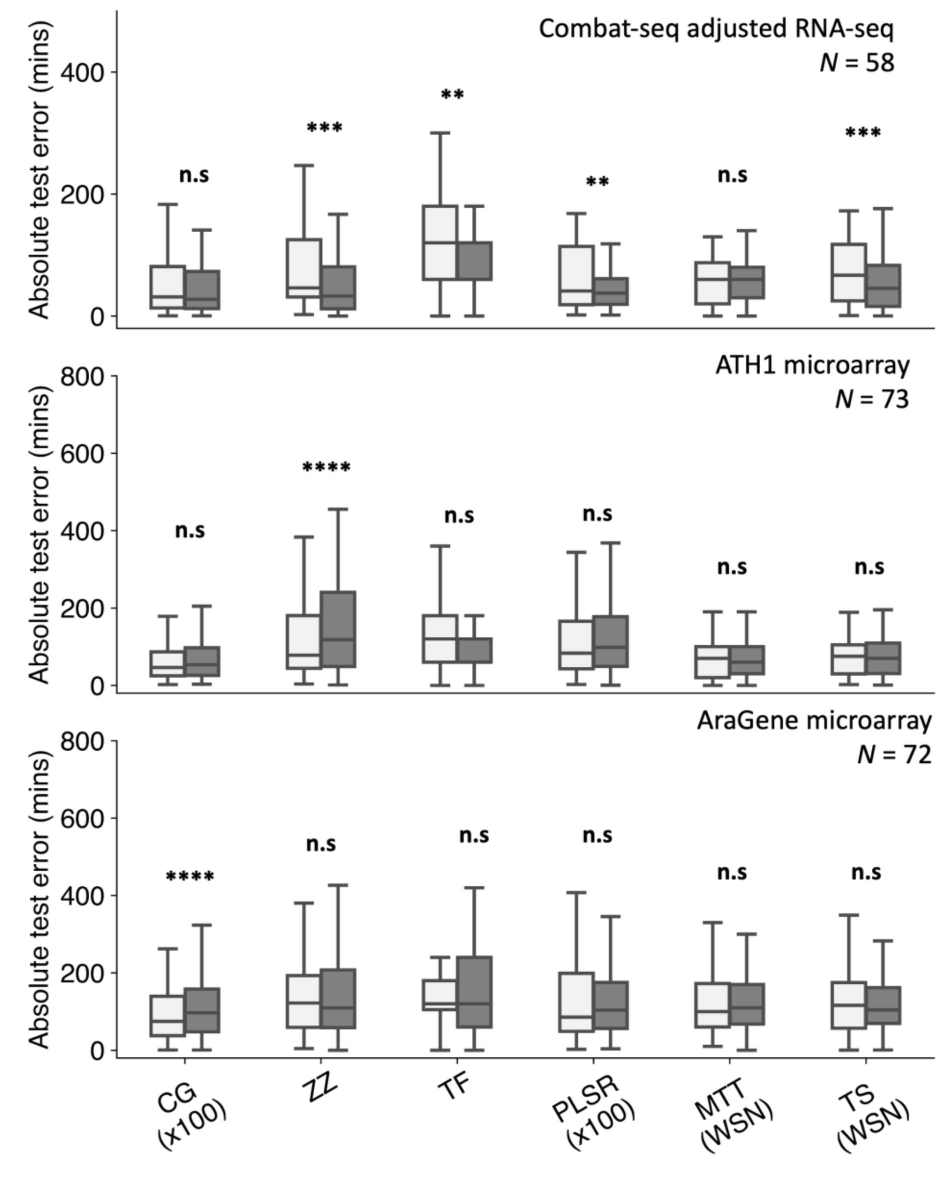


**Supplementary Figure 9:** For each circadian time (CT) prediction model, comparison of CT prediction absolute errors when fit to unadjusted and Combat-Seq[14] adjusted gene expression data. Includes ChronoGauge ensemble (CG x100), ZeitZeiger[3] (ZZ), Taufisher[5] (TF), partial-least-squares-regression[4] ensemble (PLSR x100), MolecularTimetable[6] (MTT) and TimeSignatR[7] (TS). Comparison includes a Combat-seq adjusted RNA-seq set[8–13] (*N* samples = 58), an unadjusted ATH1 microarray set[15–18] (*N* samples = 73) and an unadjusted AraGene microarray set[19] (*N* samples = 72). Methods using within-study-normalization (WSN) listed. Comparisons made using a two-tailed Wilcoxon signed-rank test, with Bonferroni adjustment of P values.

**n.s** no significance, * $P < 0.05$, ** $P < 0.01$, *** $P < 0.001$, **** $P < 0.0001$, ***** $P < 0.00001$

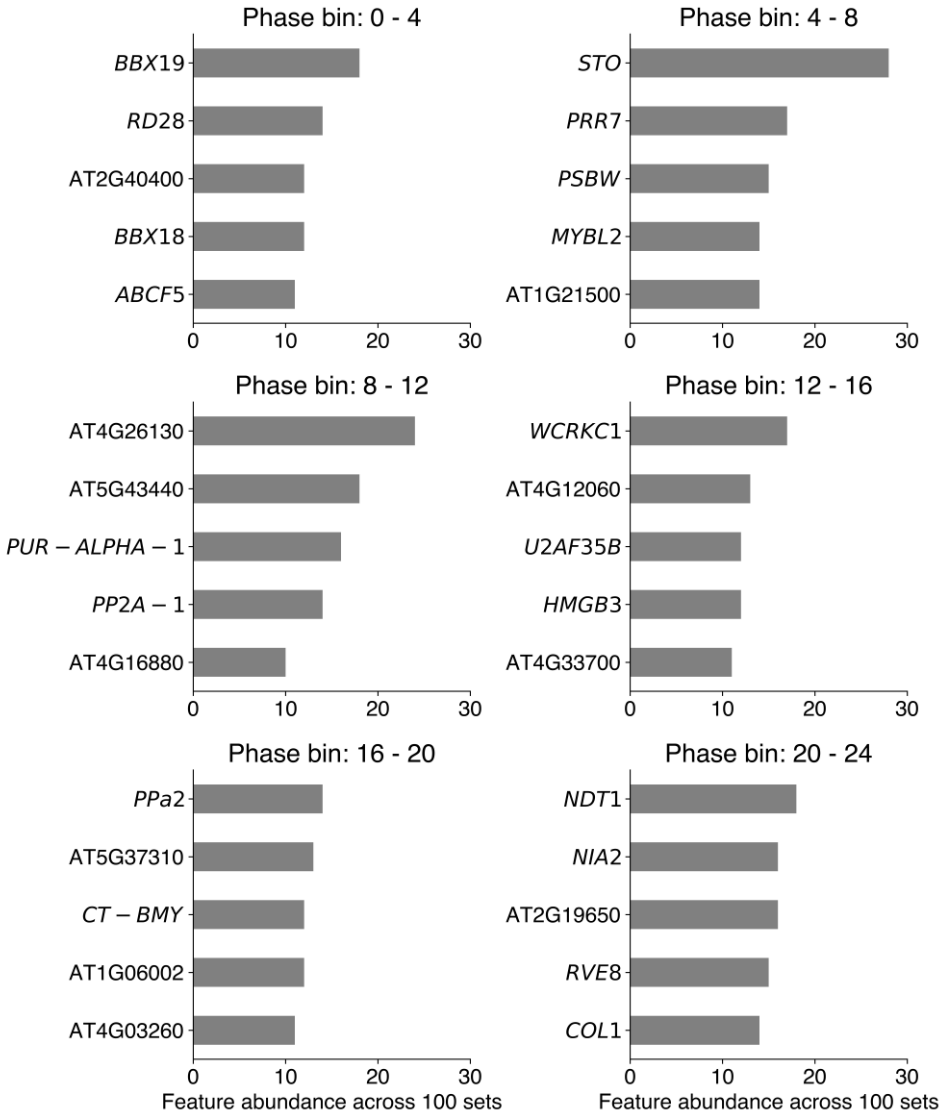


**Supplementary Figure 10:** Gene counts across the 100 feature sets of the ChronoGauge ensemble including the top 5 abundant genes from each phase bin including bins with phases ranging 0-4, 4-8, 8-12, 12-16, 16-20 and 20-24) used within the sequential feature selection (SFS) algorithm.

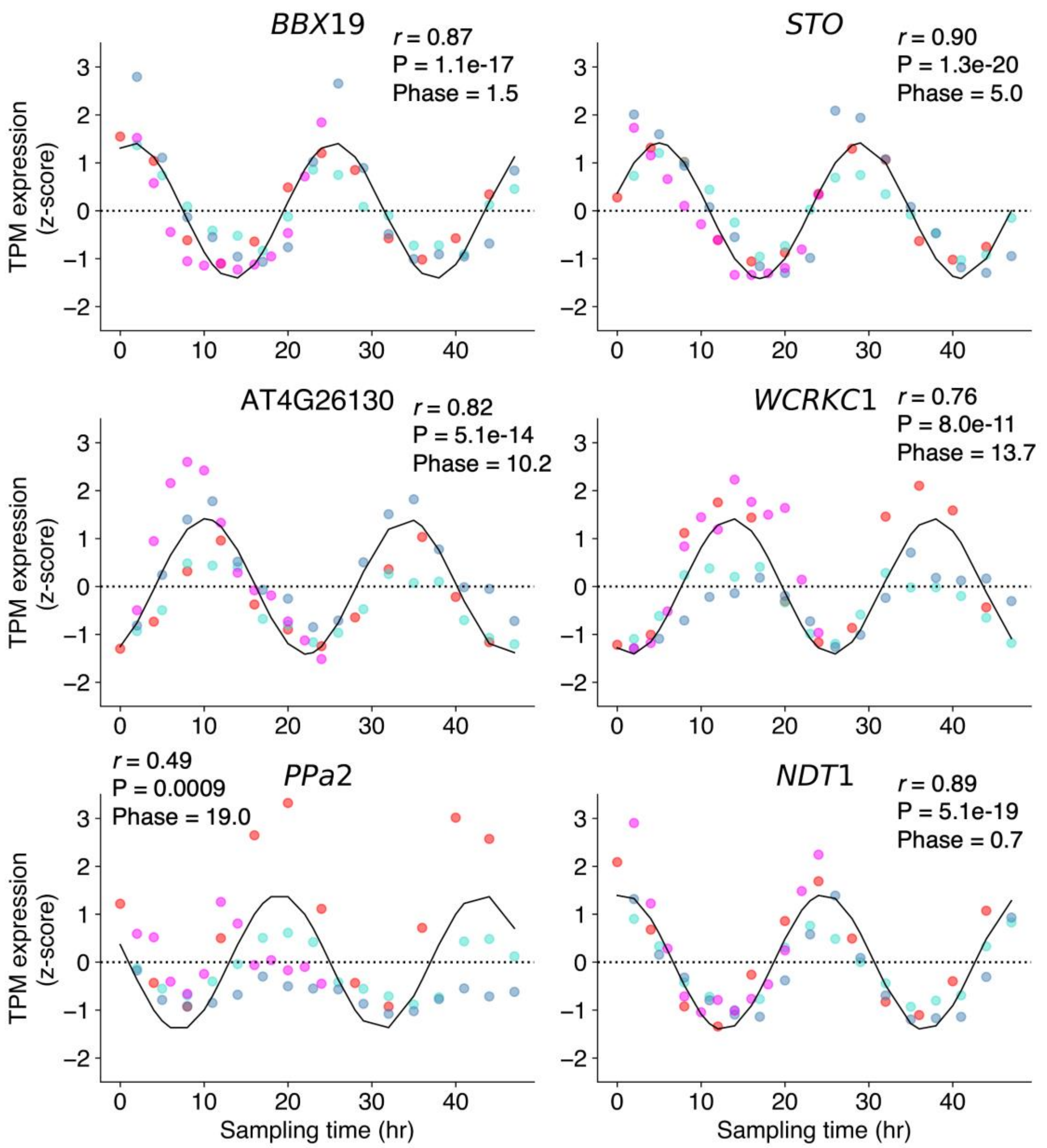


**Supplementary Figure 11:** Comparison of z-score scaled expression patterns across the training datasets[2,22,23] corresponding to the genes giving the top counts across the 100 feature sets of the ChronoGauge ensemble. Expression values included for all 4 training datasets. Curves show the optimal cosine wave fitted using the MolecularTimetable[6]-based iterative approach. Pearson correlation coefficients (*r*), Bonferroni-adjusted P-values and approximated phase listed.

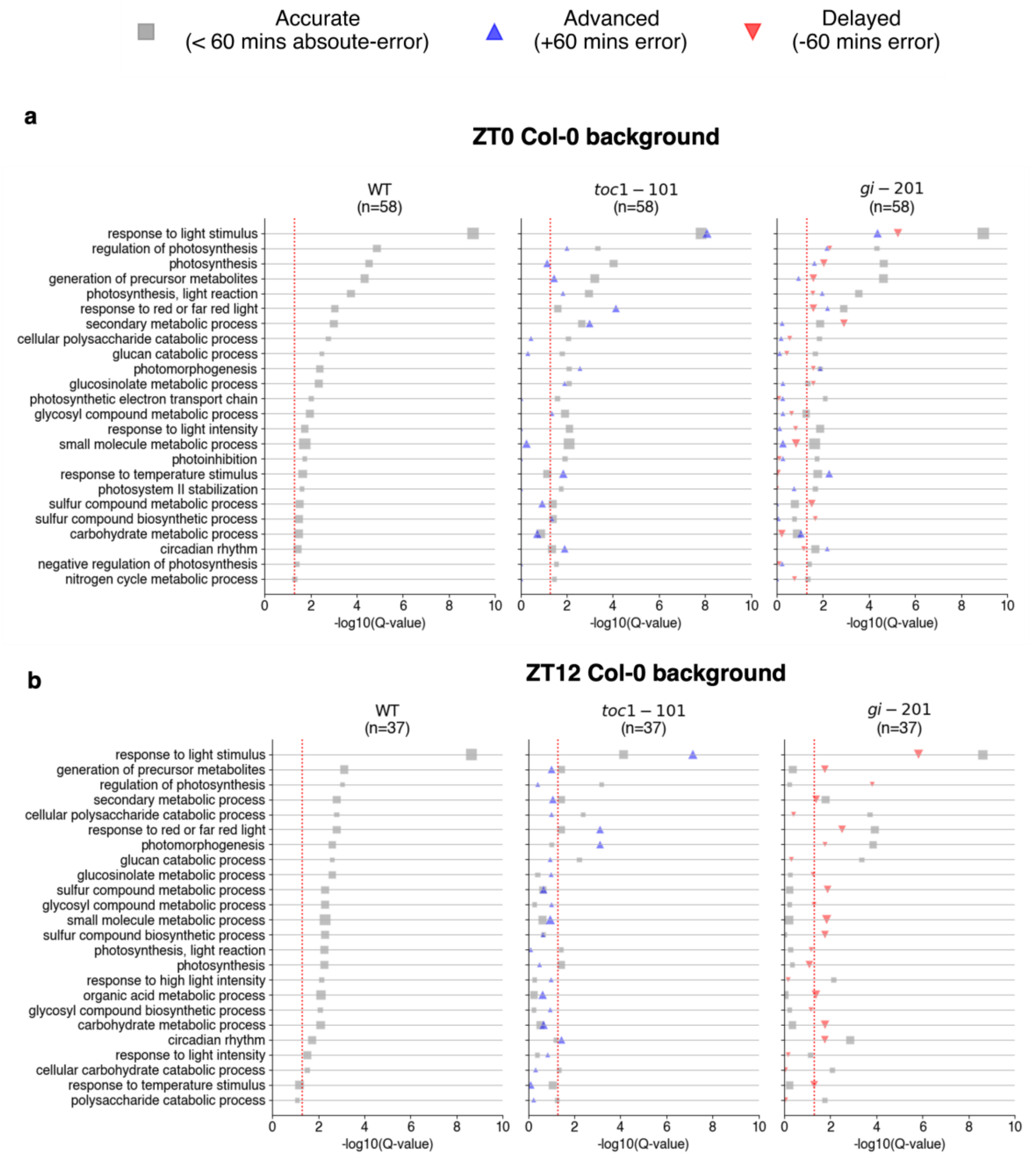


**Supplementary Figure 12:** Go term enrichment of biological processes in across wild-type (WT) and knock-out mutants *toc1-101 and gi-201* within the *Graf et al.*[12] dataset. Col-0 *Arabidopsis* samples harvested at **a** ZT0 and **b** ZT12. Significance determined using Fisher's test in TopGO[24] with Benjamini-Hochberg adjusted P values (Q values) and a threshold of Q < 0.05 (red line).

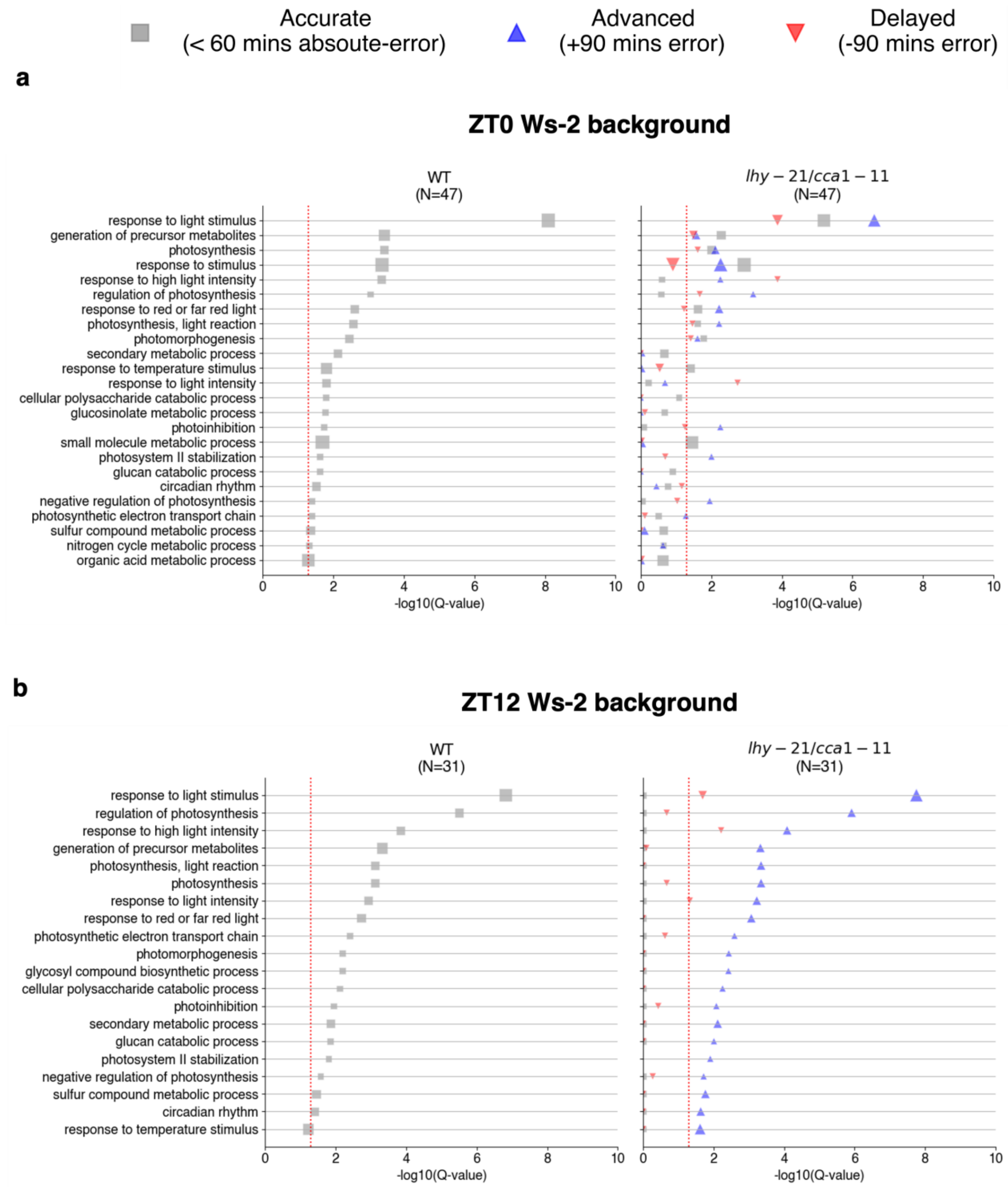


**Supplementary Figure 13:** Go term enrichment of biological processes in across wild-type (WT) plants and knock-out mutants *lhy-21/cca1-11* within the *Graf et al.*[12] dataset. Ws-2 *Arabidopsis* samples harvested at **a** ZT0 and **b** ZT12. Significance determined using Fisher's test in TopGO[24] with Benjamini-Hochberg adjusted P values (Q value) and a threshold of Q < 0.05 (red line).

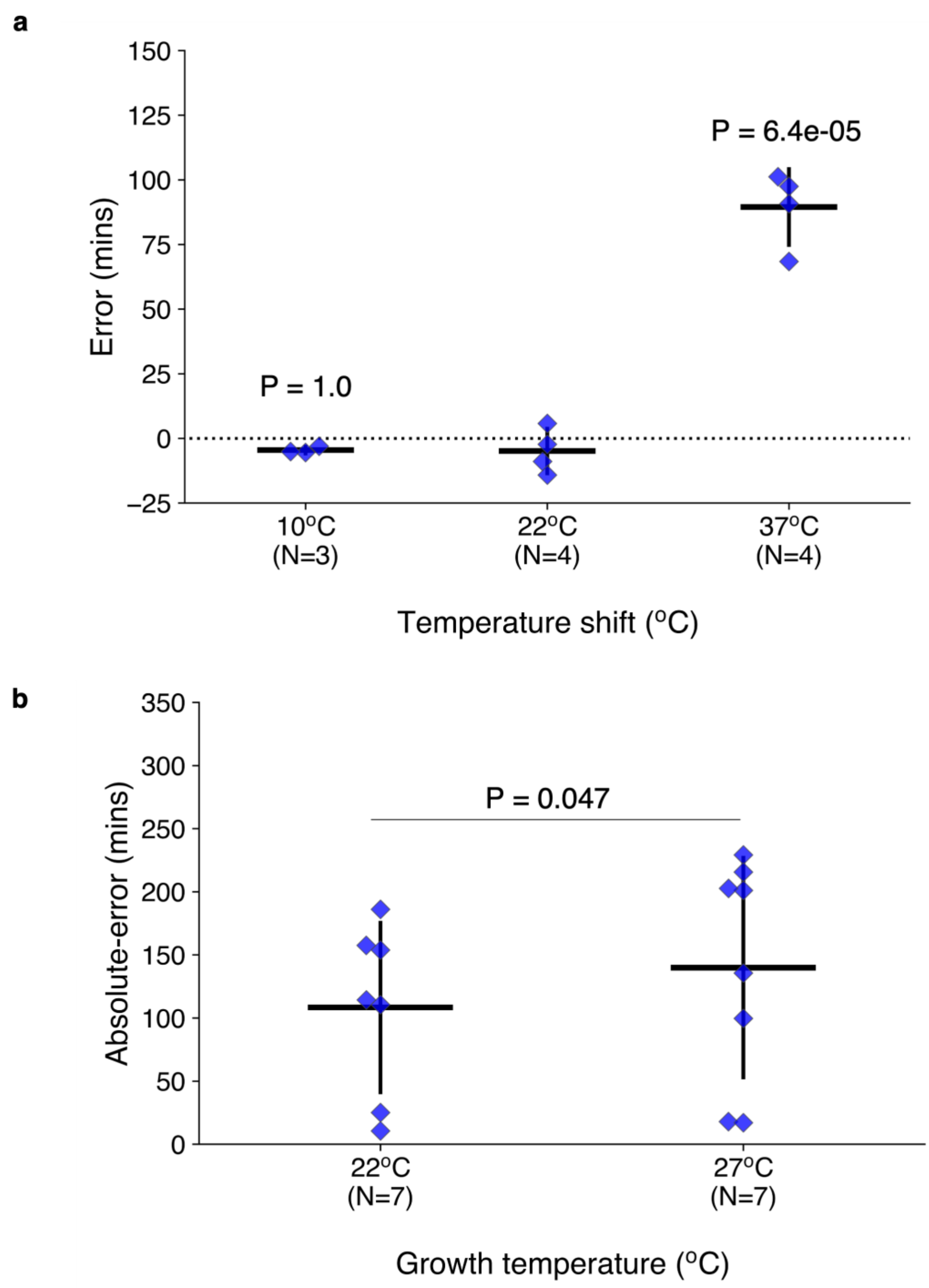


**Supplementary Figure 14:** Comparison of circadian time (CT) estimates in response to different temperature conditions. Includes **a** difference in CT errors at ZT1 after a temperature shift from 22°C to 10°C or 37°C one hour prior to sampling within *Blair et al.*[25] dataset. Significance against the control kept at 22°C determined using a two-tailed independent T-test P-values listed with correction using Bonferroni adjustment. **b** Comparison of absolute errors of CT predictions made across a time-course where plants were harvested or grown at either 22°C or 27°C within *Ezer et al.*[11] dataset. Significance determined using a Wilcoxon signed-rank test with Bonferroni adjustment of P-values.

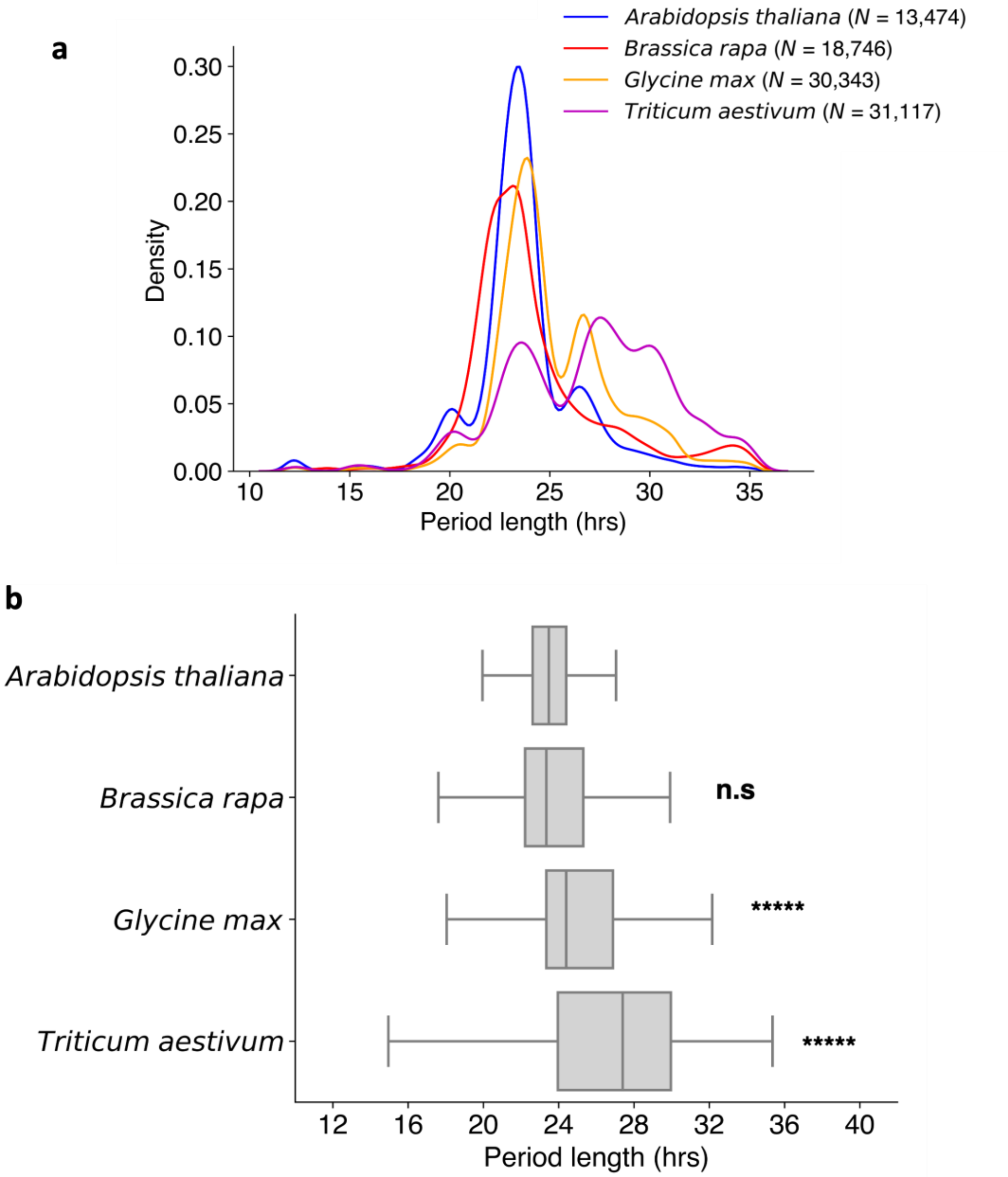


**Supplementary Figure 15:** Period length (meta2d, hrs) approximations found using Metacycle[1] for the expression of genes determined to be circadian regulated (meta2d Q < 0.05) across continuous-light (LL) time-courses for different species including *Arabidopsis thaliana* (based on the *Romanowski et al.*[2] time-course; *N* circadian regulated genes = 13,474), *Brassica rapa*[26] (*N* = 18,746), *Glycine max*[27] (*N* = 30,343) and *Triticum aestivum*[28] (*N* = 31,117). Includes **a** distribution of period lengths across each species and **b** comparison of period lengths between *A. thaliana* and other species. Comparison made using Two-tailed Mann-Whitney U test with Bonferroni adjustment of P values.

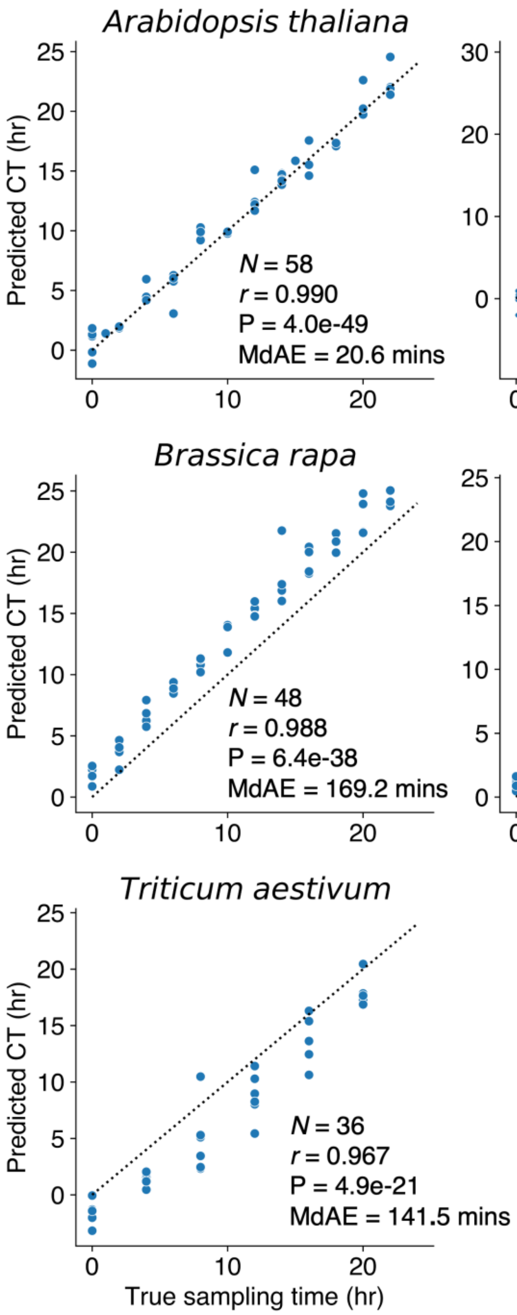


**Supplementary Figure 16:** Correlation of predicted circadian time (CT) with true sampling times across RNA-seq validation sets for *A. thaliana*[8–13] and non-model species using ChronoGauge fit only to *A. thaliana* training data[2,22,23]. For non-model species, ortholog genes were found that mapped to each of the ChronoGauge ensemble sub-predictor's *A. thaliana* gene features. Non-model species include *A. helleri*[29,30] (harvested from natural environments), *B. rapa*[26], *G. max*[27] and *T. aestivum*[28]. Pearson correlation coefficients (*r*) listed alongside Bonferroni adjusted P-values and median-absolute-errors (MdAEs).

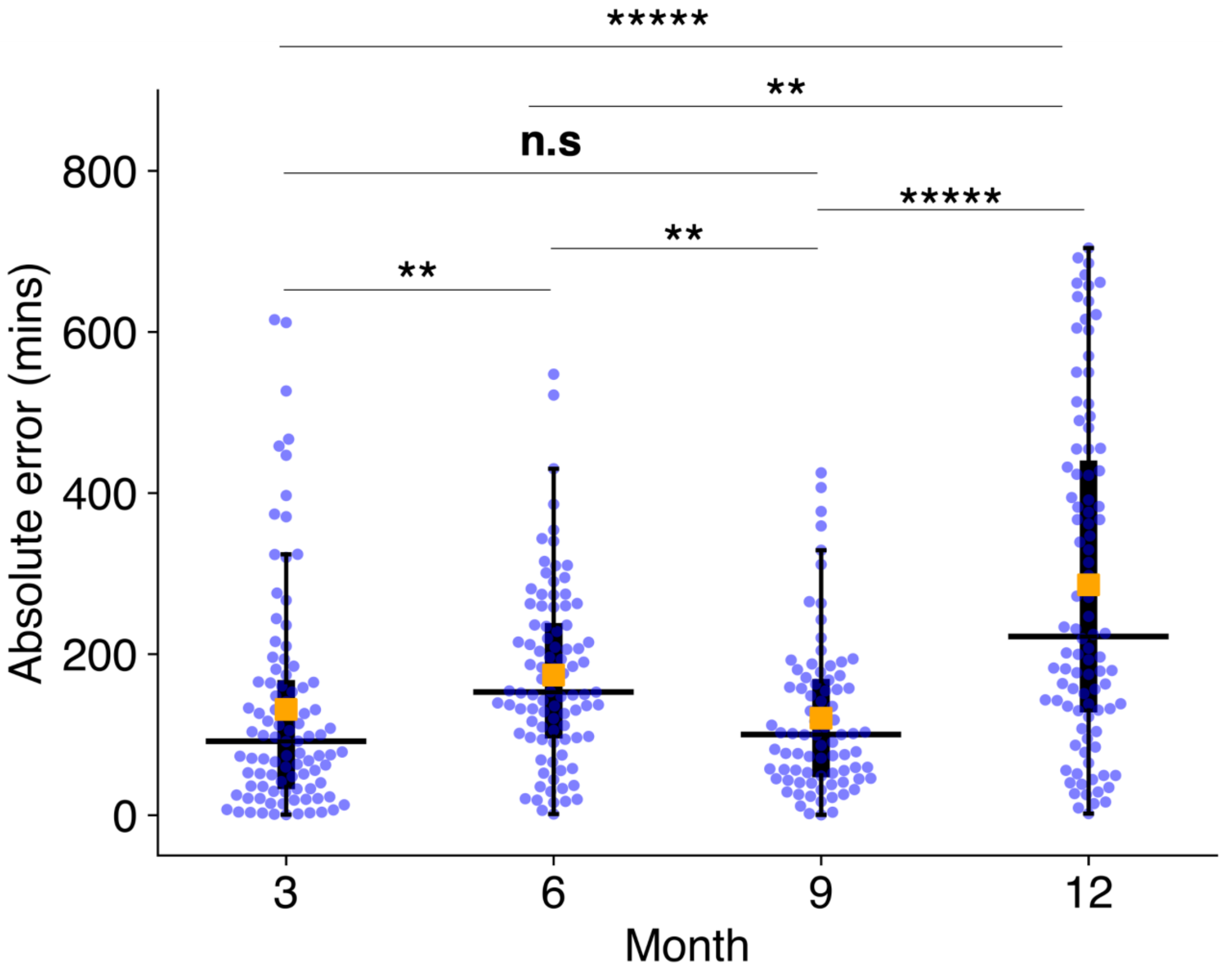


**Supplementary Figure 17:** Comparison of absolute errors of circadian time (CT) estimates made across *Arabodopsis helleri* samples from natural conditions[29,30] with associated weather data ($N = 367$). Includes median-absolute-errors (MdAEs; black line) and mean-absolute-errrors (MAEs; orange box). Absolute errors compared using a Two-tailed Mann-Whitney U test with Bonferroni adjustment of P values.

**n.s** no significance**,** ** P < 0.01**,** ***** P < 0.00001

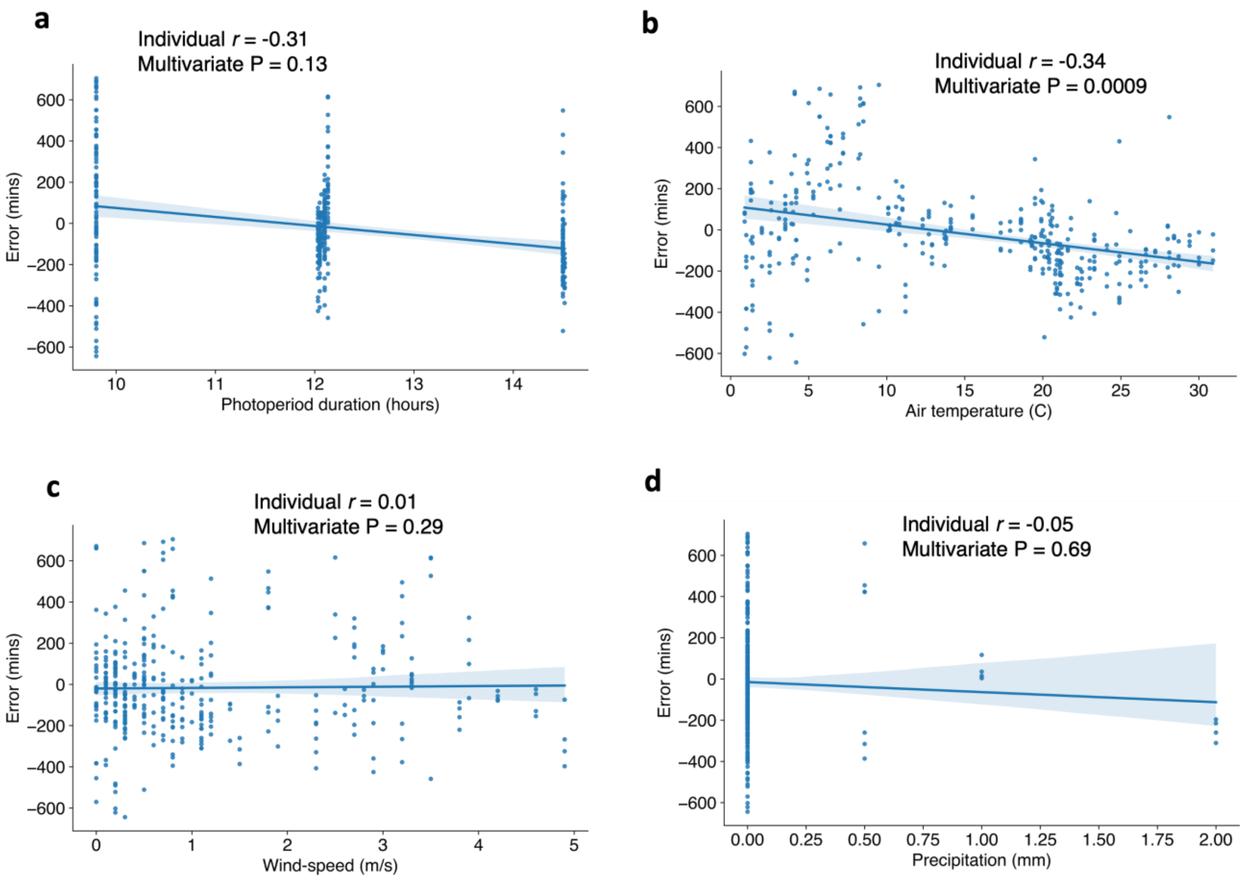


**Supplementary Figure 18:** Correlation of wild *Arabidopsis helleri* sample[29,30] circadian time (CT) prediction errors ($N = 367$) with available environmental meta-data including **a** the photoperiod duration (dusk – dawn, horurs), **b** air temperature (ºC), **c** wind-speed (m/s) and **d** the precipitation (mm) at sampling. Individual Pearson correlation coefficients (*r)* listed. Additionally, a multivariate ordinary-least-squares regression model was used to test the significance of the association while accounting for confounding environmental variables, with multivariate P values being listed.

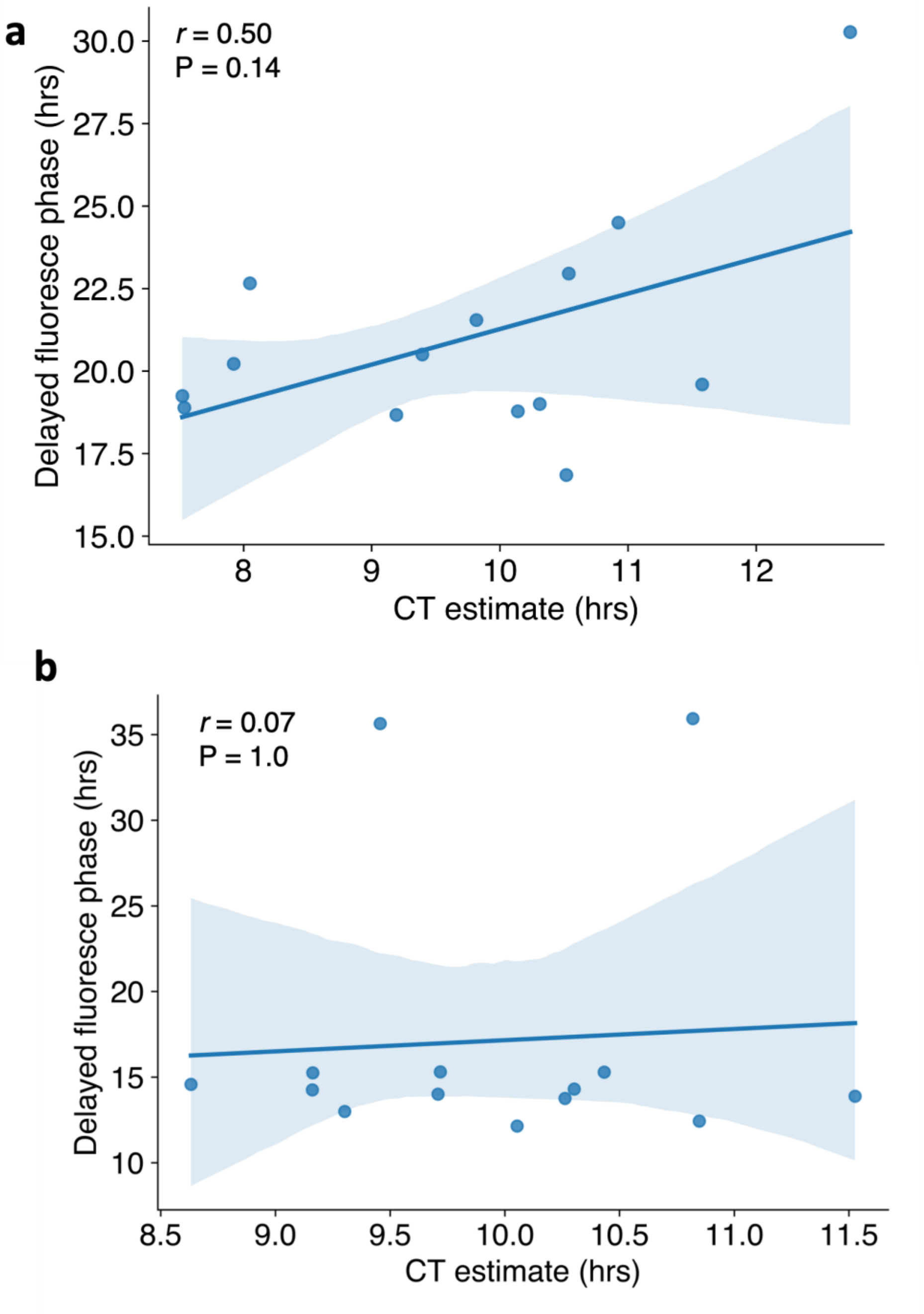


**Supplementary Figure 19:** Correlation of circadian time (CT) estimates made for *Arabidopsis* accessions ($N = 20$) using RNA-seq data by *Dubin et al*.[31] with phase approximations of the same accessions made using delayed fluorescence by *Rees et al.*[32] Includes **a** plants grown at 10ºC and **b** plants grown at 16ºC. Pearson correlation coefficient ($r$) listed with Bonferroni adjusted P values.

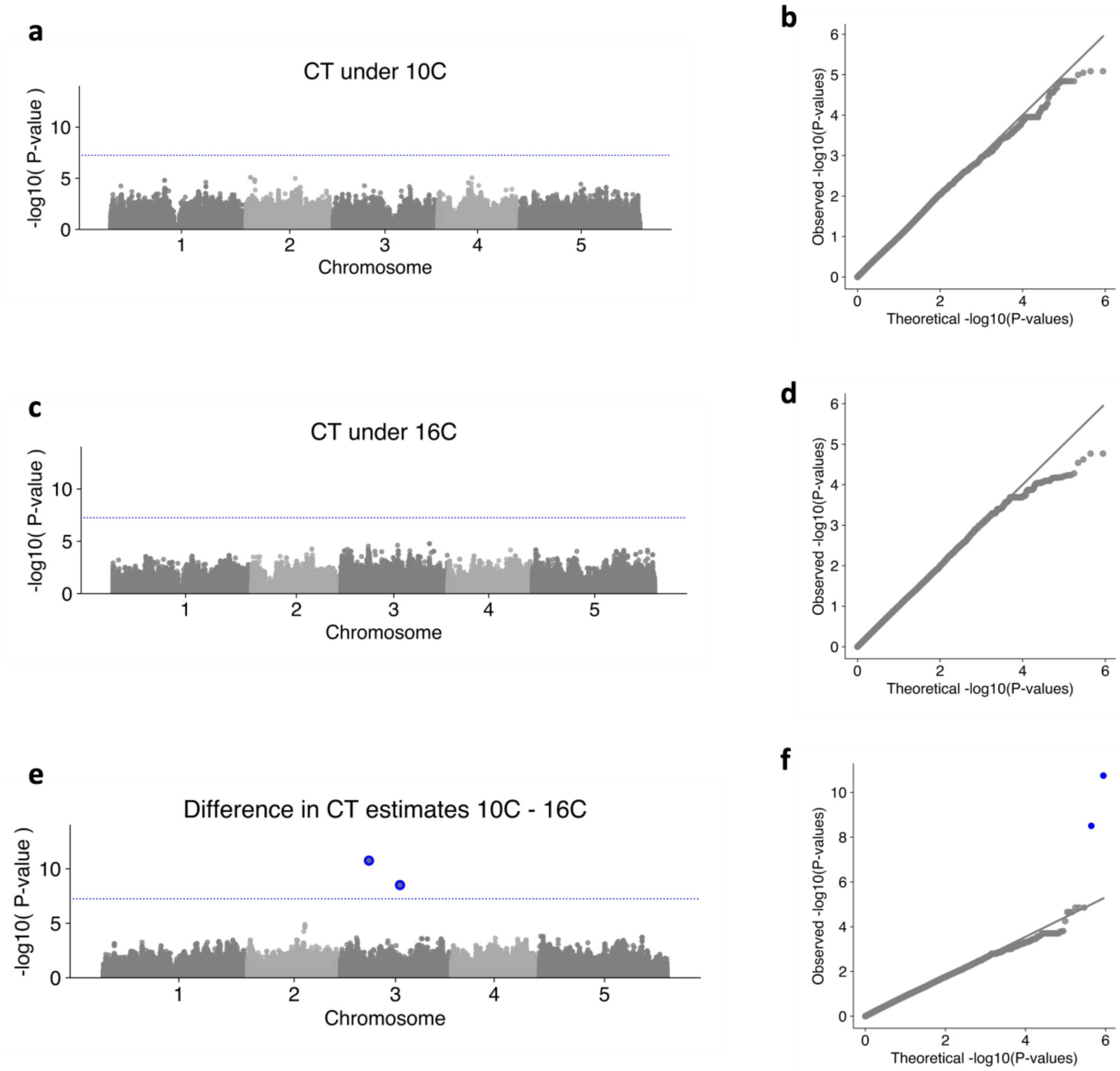


**Supplementary Figure 20:** Identification of marker-trait-associations (MTAs) across 153 *Arabidopsis* accessions within the *Dubin et al.*[31]. RNA-seq dataset using phenotypes based on circadian time (CT) estimates for each accession within the genome-wide-association-study model BLINK[33] within the R package GAPIT[34]. Genotype information for the accessions were acquired from the 1001 Genomes Project[35]. Includes significance of each single-nucleotide-polymorphism (SNP) site's association with and quantile-quantile plots using the phenotypes **a,b** CT estimates in samples grown under 10ºC, **c,d** CT estimates in samples grown under 16ºC and **e,f** difference between CT estimates grown under the two temperature groups (10ºC – 16ºC) respectively. Analyses included $N$ = 880,417 SNP sites. Significant SNPs shown (blue circle) based on Bonferroni adjustment of P values within each study. Adj. P < 0.05 corresponds to unadjusted P < 5.7e-08 (blue line).
